# Aged insulin resistant macrophages reveal dysregulated cholesterol biosynthesis, a pro-inflammatory profile and reduced foam cell formation capacity

**DOI:** 10.1101/467118

**Authors:** G Chabrier, S Hobson, N Yuldasheva, M. T Kearney, S Schurmans, I Pineda-Torra, M. C Gage

## Abstract

Insulin resistance is the central defining feature of type 2 diabetes and an independent risk factor for the development of atherosclerosis. In addition, aging is a major risk factor in the development of insulin resistance and cardiovascular disease. Macrophages play a pivotal roles the the developments of diabetes and atherosclerosis and express insulin receptors. Despite their relevance however, the effect of insulin and insulin resistance on macrophages regarding their inflammatory status, foam cell formation capacity and affect on atherosclerosis are unclear. By taking advantage of a mouse model that recapitulates the effects of chronic PI3K pathway hyperstimulation in macrophages through SHIP2 knock-down, we show that insulin resistance in aged macrophages promotes a proinflammatory phenotype while modulating the cholesterol biosynthesis pathway. These altered characteristics may contribute to the development of chronic inflammatory disease frequently observed with age. Moreover, aged insulin-resistant proinflammatory macrophages display an altered response to acute inflammatory stimulus and reduced foam cell formation capacity. Our work also highlights for the first time, the complex and contrasting immunomodulatory effects of insulin on macrophages in aged mice.

**Significance:**

- First study to explore the effects of macrophage insulin resistance in aged cells.
- Insulin resistance in aged macrophages results in the reprogramming of the transcriptome with more than 4000 genes being differentially regulated.
- Insulin resistance in aged macrophages results in upregulation of IFN signalling pathway, cholesterol biosynthesis pathway and inflammasome gene expression.
- Insulin directly regulates macrophage cholesterol biosynthesis, IFN and inflammasome gene expression in a time- and dose-dependent manner.
- Insulin resistant aged macrophages exhibit reduced capacity to become foam cells.

## INTRODUCTION

Recent changes in human lifestyle resulting in obesity and survival to an advanced age have led to a global epidemic of type 2 diabetes^1^ and cardiovascular disease^1^. Insulin resistance is the central defining feature of type 2 diabetes^2^ and an independent risk factor for the development of atherosclerosis^3^. In addition, ageing is a well known risk factor for the development of insulin resistance and cardiovascular disease^3^. The role of insulin is well established in metabolic tissues including muscle, liver and adipose tissue^4–6^ where it affects their function to regulate blood glucose levels and the term ‘insulin resistance’ is usually in reference to *metabolic* insulin resistance and the impaired ability of these metabolic tissues to respond to physiological levels of insulin. Yet insulin has significant, but less well-established roles in cells of the arterial wall^7^ and immune system including macrophages^8–20^.

Macrophages are phagocytic leukocytes of the innate immune system^21^ that play significant direct and indirect roles in the progression of diabetes and atherosclerosis through their ability to secrete a wide array of potent inflammatory cytokines, proliferate in situ^22^ and take up modified LDL in the arterial wall^23^. While the effects of insulin resistance in metabolic tissues in atherosclerosis are well defined^4–6^, the effects of insulin resistance in macrophages are less clear and often conflicting^10–12,18–20^, which may be due to a lack of consistency with the models used and end points measured. Notably, the majority of these mechanistic studies are performed exclusively on young mice, primary cells from young mice or cell lines.

Recent evidence has emerged that macrophage lipid metabolism and inflammatory response are intimately linked. For instance, down regulation of the macrophage cholesterol biosynthesis pathway can activate interferon (IFN) signalling^24^ and cholesterol accumulation can activate the inflammasome response^25^. Furthermore, a modified product of cholesterol biosynthesis downstream of the cholesterol pool, 25-hydroxycholesterol (25-OHC), has potent antiviral affects^26^ and promotes foam cell formation^27^– a critical step in the initiation and progression of atherosclerosis^28^. It is also appreciated that insulin-linked macrophage inflammation through IL-1β can acutely and directly affect systemic glucose homeostasis independently of ‘metabolic’ tissues^9^. In addition, insulin has been demonstrated to regulate cholesterol biosynthesis in liver^29^ and brain^30^.

The lipid phosphatase SHIP2 is a negative regulator of the PI(3)K signalling pathway arm of insulin^31,32^ which is activated in macrophages by insulin at physiological concentrations^9^, and it is this pathway which is also affected in insulin resistant patients^33,34^. In humans, polymorphisms of SHIP2 have been shown to be associated with obesity, diabetes and the metabolic syndrome^35–39^. Through chronic knock-down of SHIP2, we sought to investigate how chronic dysregulated insulin signalling in aged macrophages affects their inflammatory profile and interrogate their ability to form foam cells. We show that aged insulin resistant macrophages have profound changes in the macrophage transcriptome with upregulation of major inflammatory pathways including interferon signalling and inflammasome expression. In conjunction, we observe significant upregulation of the cholesterol biosynthesis pathway and the cholesterol hydroxylases *Cyp27a1* and *Ch25h* and demonstrate that aged insulin resistant proinflammatory macrophages are less able to form foam cells.

## MATERIALS AND METHODS

### Mice

Generating h-SHIP2^Δ/+^ mice. Mice were bred onto a C57BL/6J background for >10 generations in a conventional animal facility with 12-hour light/dark cycle. To examine the effect of chronically increased insulin signaling, male mice aged 50 weeks were used in all experiments, conducted in accordance with accepted standards of humane animal care under UK Home Office project license 40/3523. h-SHIP2^Δ/+^ mice were generated as in^32^. In brief: catalytically inactive SHIP2 mutant mouse was generated by inserting Cre recombinase-specific loxP sites into intronic regions flanking exons 18-19 of the *Inppl1* gene; mice with one floxed allele (SHIP2^(18-19)/+^) were crossed with *Tie2*-Cre mice (Jackson Labs) to produce progeny with germline hematopoietic-specific SHIP2 knock-down (referred to as h-SHIP2^KD^). Cre-positive SHIP2^+/-^ littermates were controls in all experiments.

### Gene expression

mRNA was isolated using TRIzol (ThermoFisher), and SHIP2 mRNA quantified using SYBR-Green based real-time quantitative PCR using (ABI Prism 7900HT, Applied Biosystems). Primer details are as follows: truncated SHIP2 forward 5’-ACC-TTA-ACT-ACC-GCT-TAG-ACA-TGG-A; truncated SHIP2 reverse 5’-ATC-AGT-GCA-ACT-AAA-TCG-AAG-GAA; non-truncated region of SHIP2 forward 5’-AAG-ACT-ACT-CGG-CGG-AAC-CA; non-truncated region of SHIP2 reverse 5’-TGCCGA-TCA-CCC-AAC-GA; β-actin forward 5’-CGT-GAA-AAG-ATG-ACC-CAG-ATC-A; β-actin reverse 5’-TGG-TAC-GAC-CAG-AGG-CAT-ACA-G.Mice were genotyped by PCR analysis of ear biopsies using Jumpstart Taq DNA Polymerase (Sigma Aldrich) and the following primers:

### Bone marrow derived macrophage culture

Bone Marrow-derived Macrophages (BMDM) were prepared as in^40^ using L929 Conditioned Medium (LCM) as a source of M-CSF for the differentiation of the macrophages. After 6 days of differentiation, LCM-containing medium was removed, cells were washed three times in warm PBS and incubated in DMEM containing low-endotoxin (≤10 EU/mL) 1% FBS and 20 µg/mL gentamycin without any LCM before being treated with insulin (time and concentrations as indicated in figure legends) or LPS (100 ng/mL, incubation times as indicated in figure legends). For LDL experiments, 5 days post-differentiation, LCM-containing medium was removed, cells washed three times with warm PBS and incubated with DMEM containing low endotoxin (≤10 EU/mL) 10% FBS and 20 µg/mL gentamycin without any LCM before being treated with native or acLDL (#5685-3404, Bio-Rad) at indicated concentrations for 24 h (mRNA quantification).

### RNA extraction and quantification

Total RNA from was extracted with TRIzol Reagent (Invitrogen). Sample concentration and purity was determined using a NanoDrop™ 1000 Spectrophotometer and cDNA was synthesized using the qScript cDNA Synthesis Kit (Quanta). Specific genes were amplified and quantified by quantitative Real Time-PCR, using the PerfeCTa SYBR Green FastMix (Quanta) on an MX3000p system (Agilent). Primer sequences are shown in supplementary table 1. The relative amount of mRNAs was calculated using the comparative Ct method and normalized to the expression of cyclophylin^41^.

### RNA array and analysis

Total RNA was extracted using TRIzol reagent (Life technologies), processed and hybridized to GeneChip™ Mouse Transcriptome Array 1.0 (MTA 1.0.) sets. Volcano plots and heatmaps were generated using Transcriptome Analysis Console Software 4.0.1 (Applied Biosystems).

### Foam cell assay and quantification

Five days post-differentiation, LCM-containing medium was removed was removed from the BMDM which were washed three times with warm PBS and incubated with DMEM containing low endotoxin (≤10 EU/mL) 10% FBS and 20 µg/mL gentamycin without any LCM and treated with native or acLDL (#5685-3404, Bio-Rad) at indicated concentrations for 24 h. Cells were then washed twice in PBS, fixed for 10 min 4% PFA, washed in PBS, rinsed in 60% isopropanol for 15 sec, incubated with Oil Red O for 1 min and rinsed for 15 sec in 60% isopropanol before 3x final PBS washes. Cells were imaged (Leica, DFC310FX) under a dissection microscope (Leica, MZ10F). Oil red O staining intensity was analysed using Image J.

### Protein isolation and immunoblotting

Total cellular protein lysates (30µg) were loaded onto a 10% SDS-PAGE gel, electrophoresed and transferred onto a PVDF membrane. The membrane was probed with anti-phosphoAkt (#4060, Cell Signaling), anti-Akt (#4691, Cell Signalling) and anti-Hsp90 (sc-7947, Santa Cruz) overnight in 2.5% BSA, TBS, followed by incubation with anti-rabbit (PO448, Dako) or anti-mouse (NA931VS, GE Healthcare) horseradish-peroxidase-tagged antibodies. Chemiluminescence (ECL 2 Western Blotting Substrate, Pierce) was used to visualise proteins.

### Statistics

Results are expressed as mean (SD). Comparisons within groups were made using paired Students t-tests and between groups using unpaired Students t tests or repeated measures ANOVA, as appropriate; where repeated *t-tests* were performed a Bonferroni correction was applied. P≤0.05 considered statistically significant.

## RESULTS

### Insulin resistance in aged macrophages results in reprogramming of inflammation and cholesterol biosynthesis gene expression

To investigate the impact of insulin resistance on macrophage inflammation in the context of aging we aged the hematopoietic SHIP2 knock-down mouse^42^ (h-SHIP2^KD^) for 50 weeks. Cultured bone marrow derived macrophages (BMDM) from these mice^40^ were compared to their Tie2Cre expressing litter mates for all experiments described. BMDM from aged h-SHIP2^KD^ mice expressed catalytically inactive SHIP2 (Fig. S1A) and were shown to be insulin resistant as assessed by the lack of phosphoAkt-S473 induction by insulin (Fig. S1B). Array profiling revealed two very different transcriptomes (Fig. S1C) showing differential expression of more than 4000 genes (2474 up- and 1673 down-regulated significantly by more than 2-fold) comparing aged h-SHIP2^KD^ to aged WT BMDM (Fig. 1A&B). Hallmark pathway analysis demonstrated that the top regulated pathways were those related to interferon immune responses (Fig. 1C, D and Fig. S1D). Upregulation of several of these known interferon responsive genes was confirmed by RT-qPCR in a separate set of experiments (Fig. S2A). The existence of an interferon-cholesterol pathway flux axis^24^ has been reported in these cells. Further interrogation of our array data revealed up-regulation of the majority of genes in the cholesterol biosynthesis pathway in the insulin resistant h-SHIP2^KD^ BMDM (Fig 1.E&F). For example, expression of *Fdps* (2.56-fold, P=0.05), *Idi1* (3.13-fold, P=0.05), *Sqle* (3.87-fold, P=0.05), *Cyp51* (4.47-fold, P=0.05), *Dhcr* (4.72-fold, P=0.05), *Msmo1* (3.25-fold, P=0.05), *Insig1* (4.06-fold, P=0.05) was observed. The differential expression of a selection of these cholesterol biosynthesis genes was subsequently confirmed by qPCR in separate experiments (Fig. S2B). In addition, cholesterol accumulation in macrophages has been linked to inflammasome activation^25^. Our array data clearly showed the up-regulation of multiple components of the inflammasome^43^ in aged h-SHIP2^KD^ BMDM compared to WT: (*Nlrp3* (1.75-fold, P=0.05), *Casp1* (2.19-fold, P=0.06), *Casp4* (4.31-fold, P=0.05) *Nlrp1b* (2.16-fold, P=0.06)), (Fig. S2C & D). Together the data suggests that insulin resistance in an aged macrophage promotes significantly increased basal inflammation potentially through modulation of cholesterol metabolism.

**Figure 1:**
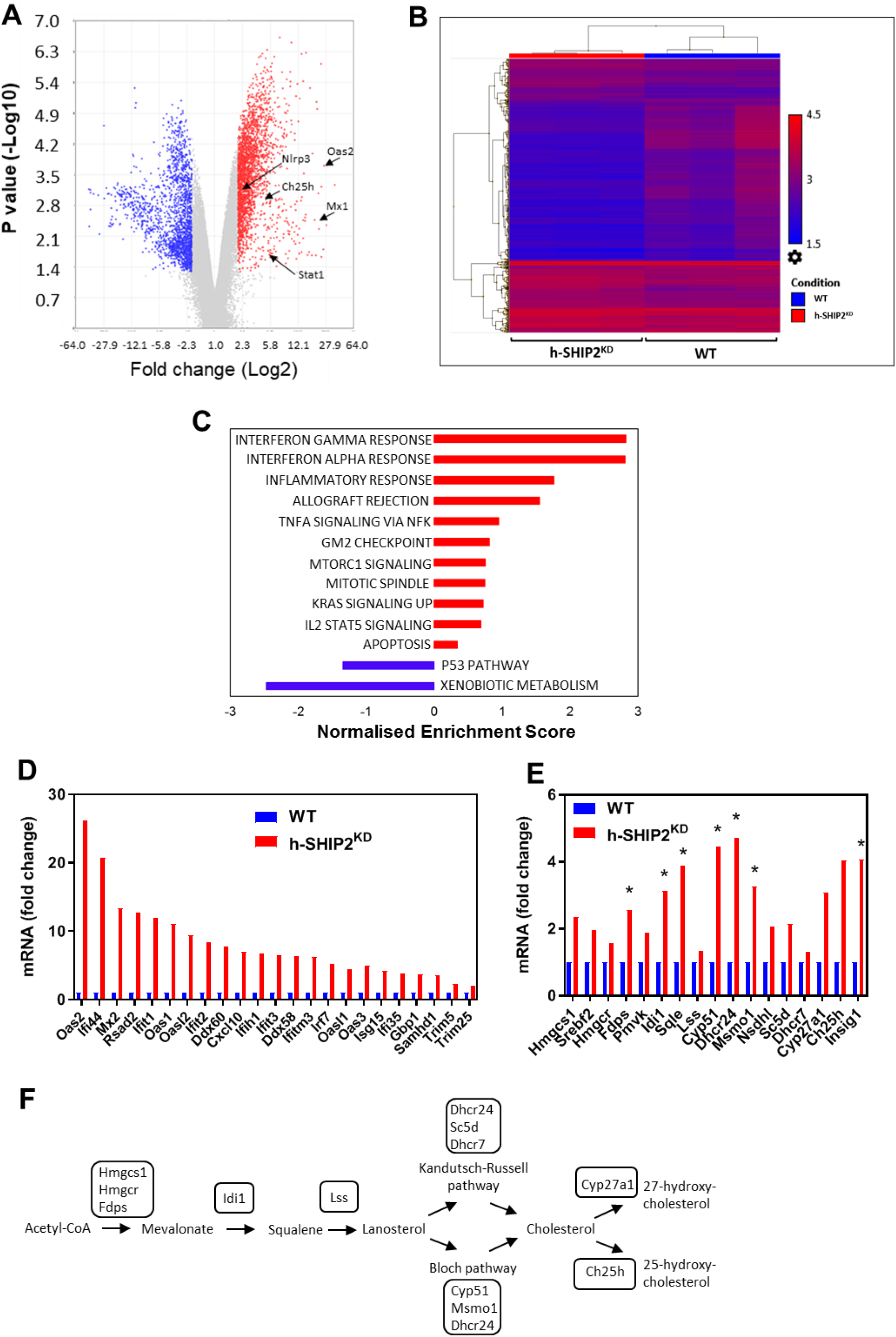
Insulin resistance in aged macrophages results in reprogramming of inflammatory profile and cholesterol biosynthesis gene expression. (**A**) Volcano plot of log2 ratio vs p-value of differentially expressed genes comparing 50 week old h-SHIP2^KD^ BMDM to 50 week old WT BMDM (n=3/group). (**B**) Clustered heatmap of RNA array gene expression (n=3 mice/group). (**C**) GSEA analysis showing enriched pathways in h-SHIP2^KD^ BMDM derived from HALLMARK gene sets. (**D**) Fold-change of interferon signature genes from array data in h-SHIP2^KD^ compared to WT (set as 1) (n=3/genotype, ≥2-fold expression, p ≤0.05 for all genes shown). (**E**) Fold-change of cholesterol biosynthesis genes in h-SHIP2^KD^ compared to WT (set as 1) (n=3/genotype, 1.5-fold expression, p ≤0.05). (**F**) Schematic of cholesterol biosynthesis pathway.

### Insulin directly regulates cholesterol biosynthesis and interferon gene expression in a time and concentration dependent manner

In normal physiology, blood insulin levels fluctuate from 0.5 -1 nmol/L when fasting, to ~5 nmol/L after a meal^44^ prior to clearance by the liver, which results in peripheral tissues receiving up to 10 fold less insulin. Insulin resistance however, results in hyperinsulinemia^45^ to compensate for reduced insulin function in metabolic tissues. Insulin has been shown to directly regulate cholesterol biosynthesis in liver^46^ and brain^30^ and insulin resistance in macrophages has been linked to cholesterol metabolism in macrophages^10,47^ although the exact mechanisms of this regulation are unknown. This could be of relevance for the development of metabolic complications in cardiovascular disease. Therefore we next investigated insulin regulation of cholesterol biosynthesis in macrophages in the context of atherosclerosis. To this end, cultured BMDM from Low density lipoprotein receptor knock-out mice (Ldlr^KO^) were exposed to insulin at a range of physiological concentrations and time points that mimic healthy, postprandial and pathophysiological levels of insulin (Fig. 2A&B). We demonstrate that BMDM cholesterol biosynthesis genes respond to acute effects of insulin (Fig. 2A&B) and interestingly the individual gene responses were dependent on concentration and duration of insulin exposure (Fig. 2A&B). For instance, 100 nM insulin for 6 hours increased *Dhcr24* mRNA expression 1.39-fold, P=0.03, but 1 nM insulin for 24 hours reduced *Dhcr24* expression 0.53-fold, P=0.01. A subset of these gene expression responses was confirmed in human primary monocytes (Fig. S3A). Insulin has been shown to be a direct inflammatory agent demonstrated through upregulation of classical proinflammatory cytokines such as TNFα^13^ and IL-1β^9^. We confirmed that insulin had an acute inflammatory effect on Ldlr^KO^ BMDM on TNFα and IL-1β expression. Notably, we found this response to be time and concentration dependent (Fig. S3B&C). We then explored whether insulin could acutely stimulate the IFN signalling genes. Consistently, separate experiments showed insulin induction of interferon genes Mx1 and Stat1 (Fig. 2C), but not other known IFN genes under the concentrations and times tested (data not shown). These IFN genes may show regulation only after longer exposure to insulin and are regulated subsequent to cholesterol biosynthesis gene regulation. This data would indicate that insulin can regulate macrophage cholesterol metabolism and inflammatory gene expression in a time and dose dependent manner.

**Figure 2:**
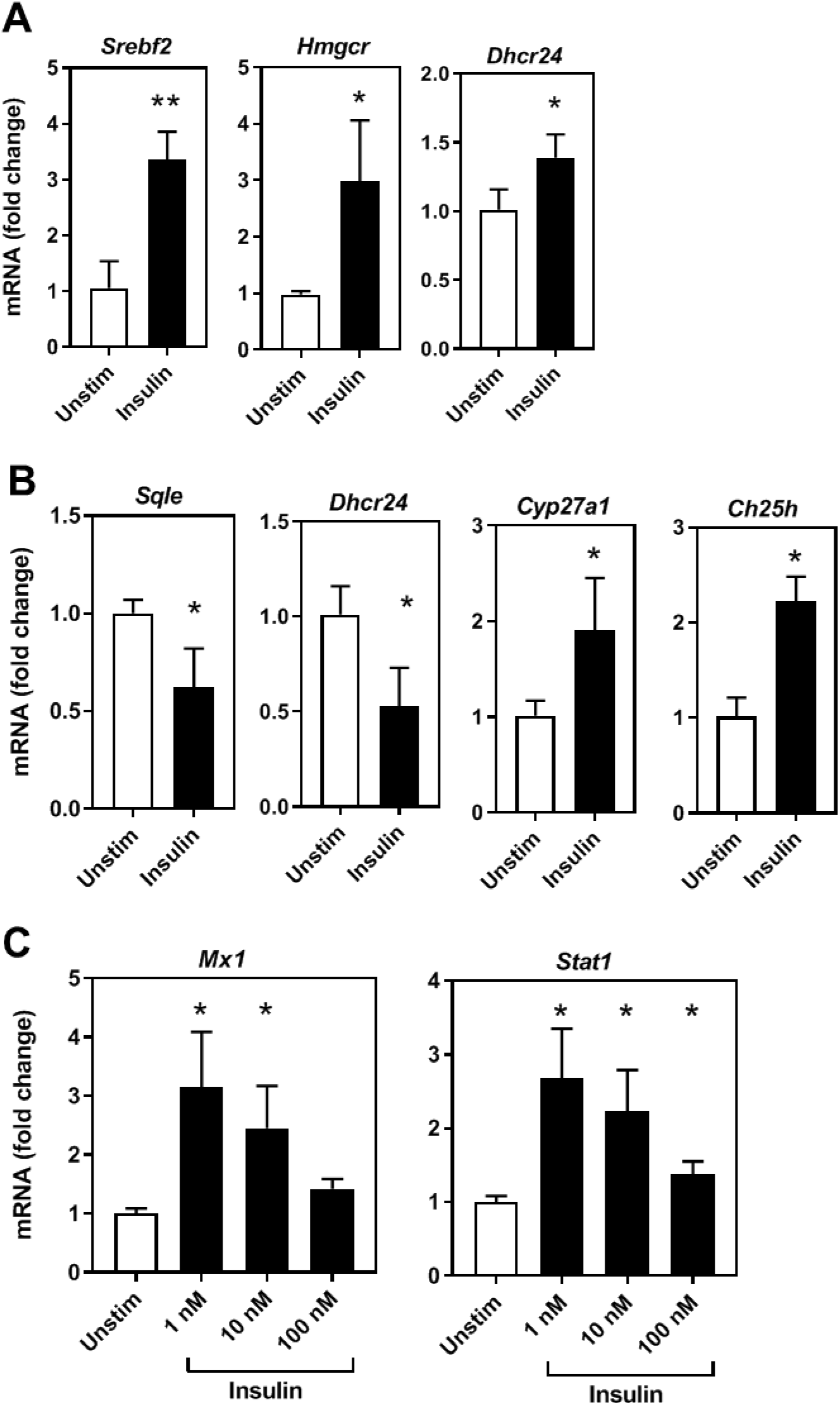
Insulin directly regulates cholesterol biosynthesis and interferon gene expression in a time and concentration dependent manner. **(A)** RT-qPCR analysis of cholesterol biosynthesis genes in Ldlr^KO^ macrophages stimulated with 100 nM insulin 6h. **(B)** RT-qPCR analysis of cholesterol biosynthesis genes in Ldlr^KO^ macrophages stimulated with 1 nM insulin 24h. **(C)** RT-qPCR analysis of interferon signalling genes in Ldlr^KO^ macrophages stimulated with insulin for 24h, representative of 3 separate experiments, p=<0.05).

### Insulin resistance in aged macrophages affects lipid metabolic and inflammatory responses to an acute LPS challenge

Macrophage inflammatory responses are linked to lipid metabolism^24,25^. As we have demonstrated that the aged insulin resistant h-SHIP2^KD^ macrophages have a dysregulated cholesterol biosynthesis and inflammatory profile (Fig. 1D&E and S2A-D), we next explored whether these insulin resistant macrophages would exhibit a differential metabolic and inflammatory response when challenged with LPS. BMDM from aged WT and h-SHIP2^KD^ mice were exposed to LPS for 6 hours. It has been shown previously that LPS-stimulated macrophages store more triglyceride and cholesterol esters and that LPS induces *Hmgcr*, the rate limiting enzyme in the cholesterol biosynthesis pathway and target of the statin family of hypolipidemic drugs^48^. We confirmed *Hmgcr* upregulation in aged WT BMDM (4.61-fold, P=0.01), and to a lesser extent in h-SHIP2^KD^ (3.58-fold, P=0.002) (Fig. 3A). Furthermore, other genes in the cholesterol biosynthesis pathway were shown to be less responsive in aged insulin resistant h-SHIP2^KD^ macrophages (Fig. 3A). LPS has also be observed to be a strong inducer of the cholesterol hydroxylase *Ch25h*^49^ which converts cholesterol to 25-hydroxycholesterol (25-OHC). Beyond its metabolic role in cholesterol homeostasis as a negative regulator of cholesterol biosynthesis^26^, 25-OHC has been shown to have important direct antiviral functions^26^, modulate adaptive immune responses^49^ and affect foam cell formation through lipid droplet retention^27^. In addition to *Hmgcr*, expression of *Ch25h* was also shown to be less responsive to acute LPS challenge in the aged h-SHIP2^KD^ cells compared to aged WT control BMDM (*Ch25h* 68-fold, P=<0.0001 vs 189-fold, P=0.0002 respectively) (Fig. 3A, right panel). In keeping with this less responsive phenotype, the levels of IFN signaling genes were also shown to be less responsive to LPS in h-SHIP2^KD^ macrophages (Fig. 3B). Intriguingly, components associated with the inflammasome showed mixed responses – Nlrp3 and caspases 1 and 4 showed decreased responses (*Nlrp3* 67-fold, P=<0.0001 vs 122-fold, P=<0.0001, *Casp1* 2.87-fold, P=0.0004 vs 5.65-fold, P=<0.0006, *Casp4* 5.97-fold, P=0.0002 vs 23-fold, in mutant and WT macrophages respectively) (Fig. 3C). By contrast, mRNA expression of cytokines know to be influenced by the NLRP3 inflammasome IL-18 and IL-1β showed an increased response to LPS in the mutant cells (*Il-18* (20.53-fold, P=<0.0001 vs 11.13-fold, P=0.0002), (*Il-1β* (395-fold, P=<0.0001 vs 261-fold, P=0.0001) (Fig. 3C).

**Figure 3:**
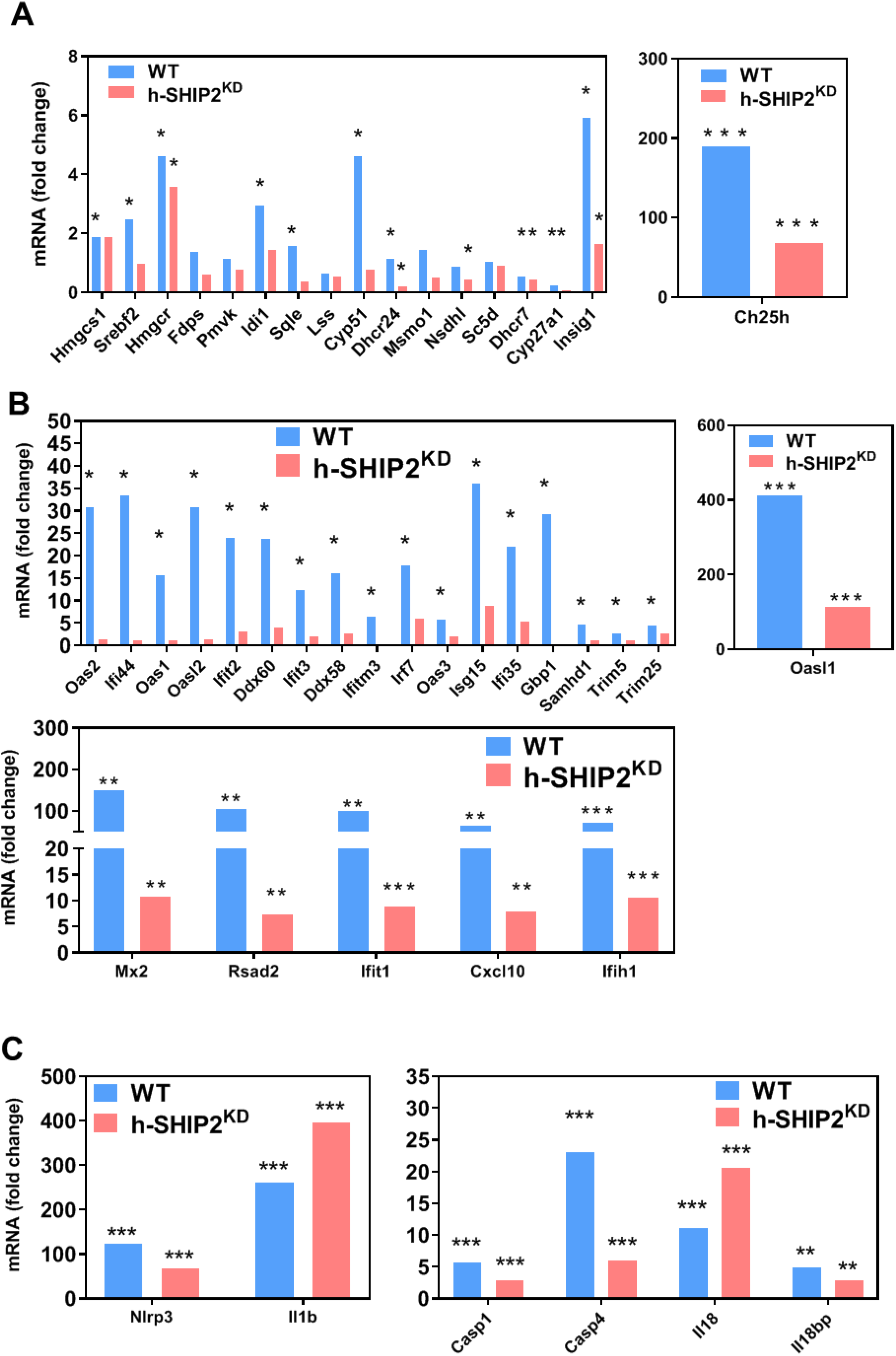
Insulin resistance in aged macrophages affects lipid metabolic and inflammatory responses to acute stimulus. (**A**) Fold-change of cholesterol biosynthesis genes from LPS 6h array data in aged h-SHIP2^KD^ and WT BMDM compared to basal (set as 1) (n=3/genotype, ≥2-fold expression, p ≤0.05). (**B**) Fold-change of interferon signature genes from LPS 6h array data in aged h-SHIP2^KD^ and WT BMDM compared to basal (set as 1) (n=3/genotype, ≥2-fold expression, p ≤0.05). (**C**) Fold-change of inflammasome associated genes from LPS 6h array data in aged h-SHIP2^KD^ and WT BMDM compared to basal (set as 1) (n=3/genotype, ≥2-fold expression, p ≤0.05).

As our insulin stimulation data have highlighted the important roles that time and concentration of insulin play in macrophage cholesterol metabolism and inflammatory responses (Fig. 2 and Fig. S3), we also investigated LPS responses in aged BMDM after a longer 24 hour incubation period. Consistent with our previous observations at 6h (Fig. 3A), *Hmgcr* expression showed significant up-regulation in WT and h-SHIP2^KD^ BMDM after 24 hrs of LPS (*Hmgcr* 5.54-fold, P=<0.0008 vs 5.29-fold, P=0.02 respectively) (Fig. S4A) as did further genes in the cholesterol biosynthetic pathway (data not shown). However, in striking difference to the short term LPS challenge, genes involved in cholesterol catabolism including *Ch25h* were negatively regulated after 24hrs of LPS challenge in WT and h-SHIP2^KD^ (*Ch25h* 0.16-fold, P=0.002 vs 0.27-fold, P=0.02 respectively) (Fig. S4B). Recently it has been shown that at high concentrations of insulin, components of the inflammasome including *Nlrp3* are up-regulated in M1 classically-activated macrophages but not in unstimulted or MO macrophages^9^. Therefore we next explored if the aged insulin-resistant pro-inflammatory h-SHIP2^KD^ macrophages would also demonstrate inflammasome upregulation in response to insulin (Fig. S4D). We observed no induction in WT (MO) macrophages, intriguingly however, we observed a trend towards negative regulation of *Nlrp3*, *Il-1β*, *Il-18* and *Il-18bp* in h-SHIP2^KD^ (Fig. S4D). Expression of inflammasome components in response to LPS challenge was shown to be significantly increased in the h-SHIP2^KD^ compared to WT (Fig. S4E) however we observed a repeated trend towards anti-inflammatory actions of insulin when co-incubated with LPS (Fig. S4E). Taken togther, the data points to aged insulin resistant macrophages exhibiting increased inflammation at a basal level, and are generally less able to respond to acute inflammatory challenge.

### Insulin resistance in aged macrophages reduces foam cell formation capacity and alters Ch25h and inflammasome response to modified LDL

Insulin resistance is an independent risk factor for atherosclerosis^3^. We have observed differences in cholesterol metabolism including cholesterol hydroxylase enzyme expression *Ch25h* shown to play a role in foam cell formation through lipid droplet retention. We investigated how insulin resistance in aged macrophages would affect foam cell formation capability. Aged WT and aged h-SHIP2^KD^ macrophages were incubated with acetylated LDL or native LDL for 24 hours. We found that aged h-SHIP2^KD^ BMDM showed resistance to foam cell formation (Fig. 4A) when compared to aged WT controls. We next explored changes in mRNA expression of cholesterol biosynthesis metabolism genes in these foam cells. As observed by others^50^, expression of genes in this pathway were negatively regulated in response to cholesterol loading, however these responses were similar between h-SHIP2 and WT (Fig. S5A). In contrast, we observed differences in the response to acLDL in the cholesterol hydroxylase genes *Cyp27a1* and *Ch25h* when comparing WT to h-SHIP2^KD^ cells (*Cyp27a1* 0.87-fold, P=0.16 vs 0.46-fold, P=0.025, *Ch25h* 0.98-fold, P=0.75 vs 0.39-fold, P=0.003 respectively) (Fig. 4B). Cholesterol accumulation has been linked to activation of the inflammasome in young mouse BMDM^25^. Thus, we next investigated inflammasome gene expression response to acLDL loading in h-SHIP2^KD^ vs WT BMDM. We found that *Casp1*, *Il-1β* or *Il-18* responses was not affected by modification of insulin signalling through the PI3K pathway (Fig. S5B). Interestingly though, *Nlrp3* was down-regulated in h-SHIP2^KD^ vs WT BMDM (*Nlrp3* 0.49-fold, P=0.06 vs 0.8-fold, P=0.12, respectively) (Fig. 4C) which may indicate a difference between young and old insulin resistant macrophages response to cholesterol loading. The data show that aged insulin resistant macrophages have reduced ability to form foam cells, and that this may be due to increased activity of cholesterol hydroxylases *Cyp27a1* and *Ch25h* modulating intracellular cholesterol levels.

**Figure 4:**
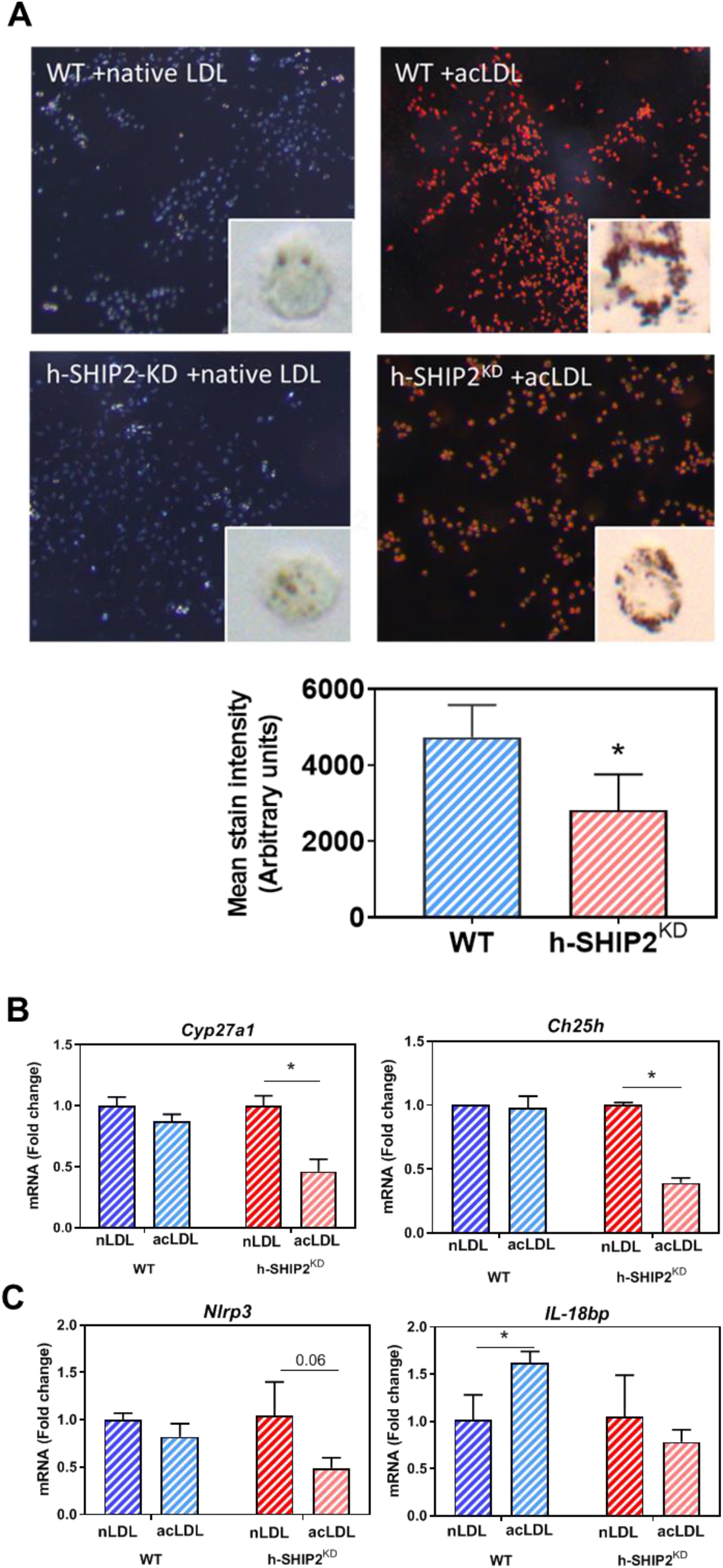
Insulin resistance in aged macrophages reduces foam cell formation capacity and alters Ch25h and inflammasome response to modified LDL. **(A)** h-SHIP2^KD^ BMDM are protected against foam cell formation compared to WT (incubated with 50ug native/acetylated LDL 24hrs, stained with Oil Red O, main image x80 mag, inset image: single cells showing uptake of lipid droplets x200 mag, analysis of 3 images, triplicate samples, n=3). **(B)** RT-qPCR analysis of cholesterol hydroxylase genes in aged h-SHIP2^KD^ and WT macrophages stimulated with 50ug native/acetylated LDL 24hrs. **(C)** RT-qPCR analysis of inflammasome associated genes in aged h-SHIP2^KD^ and WT macrophages stimulated with 50ug native/acetylated LDL 24hrs.

## DISCUSSION

The effects of macrophage insulin resistance and inflammation in the context of atherosclerosis are unclear. Indeed several studies demonstrate that macrophage insulin resistance promotes atherosclerosis or foam cell formation^10,14,19^ while a similar number of reports show that macrophage insulin resistance is protective^11,17,20^. These discrepancies may be due to the different models of insulin used and time points explored. Remarkably, while ageing is a significant risk factor for insulin resistance and atherosclerosis, the majority of these studies are performed on young mice^51^, cells from young mice, or cell lines. Furthermore, the normal physiological role of insulin on macrophages is yet to be fully appreciated as several studies have shown insulin to have anti-inflammatory functions^8,15^, while others demonstrate pro-inflammatory functions of insulin^9,13,18,52^ on macrophages. Here we sought to determine the effects of insulin resistance in aged macrophages on inflammatory phenotype and their capacity to form foam cells – a pivotal step in the progression of atherosclerosis.

### Chronic upregulation of the PI3K pathway in macrophages through SHIP2 knock-down results in macrophage insulin resistance

While previous models have employed insulin receptor ablation or knock-out of other components of the insulin signalling pathway to study insulin resistance, we utilised BMDMs with chronic SHIP2 knock-down (h-SHIP2^KD^) from our recently generated mouse model of aged insulin resistance^42^. SHIP2 is a negative regulator of the PI3K signalling pathway arm of insulin^31,32^ and it is this pathway which is activated in macrophages by insulin at physiological concentrations^9^, and it is this pathway which is affected in insulin resistant patients^33,34^. Knocking-down SHIP2 mimics hyperstimulation of this pathway as seen in insulin resistant hyper-insulinemic patients^34^. We show that chronic SHIP2 knock-down in macrophages from 50 week-old mice results in insulin resistance (Fig. S1B).

### Aged insulin resistance in macrophages reprograms the transcriptome up-regulating proinflammatory pathways and cholesterol biosynthesis enzyme expression

Transcriptome array analysis reveal more than 4000 genes which were differentially expressed >2-fold with major upregulation of the interferon signalling pathways (Fig. 1C&D). Recent work has demonstrated there exists a crucial feedback loop between macrophage cholesterol pathway flux and interferon signalling^24^. Our data also shows upregulation of cholesterol biosynthesis enzyme genes (Fig. 1E&F and S2B) as observed in cholesterol starved cells. Insulin has been demonstrated to directly regulate cholesterol biosynthesis in liver^29^ and brain^30^, and our novel findings now suggest that insulin resistance in macrophages dysregulates this pathway in aged macrophages.

### Insulin can directly activate cholesterol biosynthesis and interferon genes in macrophages

As insulin has been demonstrated to directly regulate cholesterol biosynthesis in liver^29^ and brain^30^ we sought to determine if insulin can directly regulate genes of the cholesterol biosynthetic pathway in BMDM. We observed that insulin can directly up- or down-regulate these genes in a time and concentration dependent manner (Fig. 2A&B and S3A) which is befitting of a hormone which itself is cyclic/phasic in its behaviour in both normal physiology and insulin resistance pathology. As others^9,13^, we observed that insulin can act directly as a proinflammatory agent in macrophages (Fig. S3B&C), and we went on to show that a limited number of interferon genes were consistently activated by insulin (Fig.2C). However, other IFN genes were not, as observed from the chronic dysregulated insulin-signalling array data (Fig.1D), which may reflect the significance of exposure times to insulin that macrophages require for dysregulation in a pathophysiological setting.

### Aged insulin resistant proinflammatory macrophages respond acutely to LPS through further up-regulation of Il-1β and Il-18

We found that upon acute exposure to LPS, aged WT macrophages responded in a typical fashion through upregulation of cholesterol biosynthesis pathway genes (Fig.3A), interferon signature genes (Fig.3B) and inflammasome associated genes (Fig.3C). However, h-SHIP2^KD^ BMDM reacting from a higher pro-inflammatory basal state (Fig. 1C &D and S2A,C-D) were less able to respond to acute LPS challenge except for significantly raised expression of *Il-1β* and *Il-18* (Fig.3C). Taken together, the data would suggest that insulin resistance promotes a proinflammatory macrophage phenotype with atered potential to respond to acute inflammatory stimulus such as that seen in infection.

### Insulin has an anti-inflammatory effect

While we show that insulin has a direct inflammatory effect on macrophages (Fig.2C and S3B&C, we also observed anti-inflammatory properties of insulin when incubated with LPS across a range of inflammasome associated (Fig.S4D) and IFN signature genes (Fig.S4E). Previous reports have demonstrated anti-inflammatory action of insulin on macrophages^8,15^. Overall, our data indicate that insulin may have a dual role in inflammation, perhaps acting as a pro-inflammatory agent in an unactivated (MO) macrophage, and an anti-inflammatory agent in an activated macrophage.

### Aged insulin resistant proinflammatory macrophages are less able to form foam cells

We show that upon loading with modified LDL, aged macrophages with defective insulin signalling are less able to form foam cells (Fig.4A) – which is a pivotal step in the progression of atherosclerosis. Interestingly, we find changes in responses of the cholesterol hydroxylase genes downstream of the cholesterol pool: *Cyp27a1* and *Ch25h* (Fig. 4B). A ground-breaking report has recently shown that non-foamy, rather than foamy macrophages, have a proinflammatory profile^53^ and that peritoneal macrophages when challenged with modified LDL accumulate desmosterol (a cholesterol biosynthetic intermediate) which suppressed inflammatory gene expression)^50^. Furthermore, 25-hydroxycholesterol (the product of *Ch25h*) also has direct role in inflammation^26^ and foam cell formation^27^. Our work therefore extends previous reports that macrophage insulin resistance is protective against foam cell formation^11,12^ to an aging setting and suggests that the role of Ch25h may play a significant role in this process.

Collectively, this study demonstrates that in a physiologically relevant model of aged insulin resistance (h-SHIP2^KD^), macrophages display a proinflammatory transcriptome which may contribute to the development of chronic inflammatory disease frequently observed with age. This heightened inflammatory state responds abnormally to acute inflammatory stimulus and compromises the macrophages ability to form foam cells (Fig. 5 A-C). Our work also highlights for the first time the complex and contrasting immunomodulatory effects of insulin on macrophages in aged mice.

**Figure 5:**
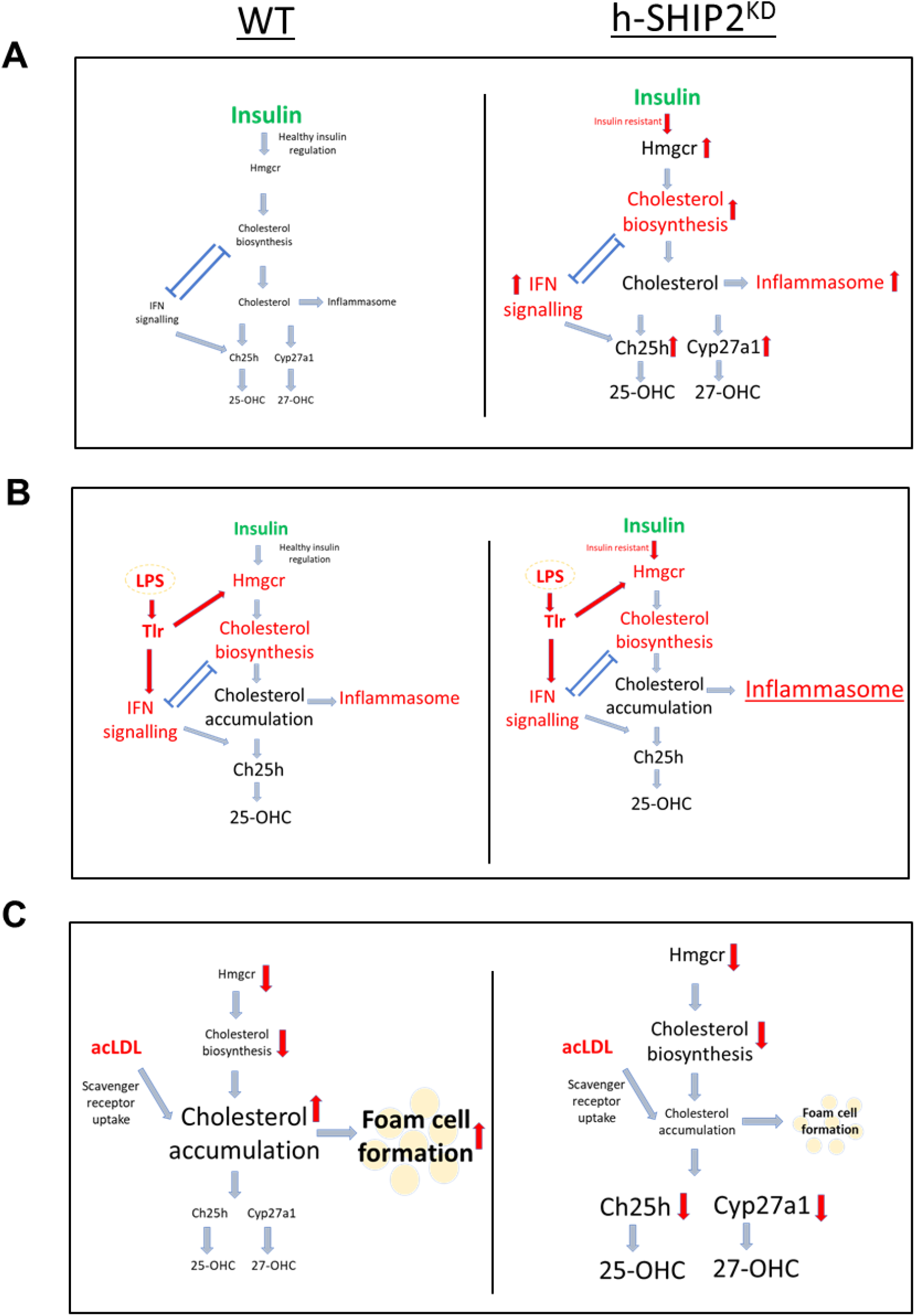
Summary schematic. **(A)** Insulin resistance in aged macrophages results in proinflammatory phenotype, which **(B)** demonstrates selective inflammatory response to acute inflammatory stimulus and **(C)** are less able to form foam cells.

## ACKNOWLEDGEMENTS

This work was supported by a Diabetes UK Small Project Grant 15/0005195 (MG), British Heart Foundation Project Grant PG/16/87/32492 (MG) and a Diabetes UK Project Grant 17/0005682 (MG).

## AUTHOR CONTRIBUTIONS

M.G conceived the study, secured funding, designed experiments, performed experiments, analysed and interpreted data, wrote the manuscript and supervised all aspects of the work. G.C performed experiments and analysed data. S.H performed experiments and analysed data. N.Y collected and supplied samples. M.K provided samples, helped secure funding. S.S generated SHIP2floxed mouse^32^, IPT helped secure funding and provided critical revision of manuscript.

## REFERENCES

1. King, H., Aubert, R. E. & Herman, W. H. Global burden of diabetes, 1995-2025: prevalence, numerical estimates, and projections. Diabetes Care 21, 1414–31 (1998).

2. Benito, M. Tissue-specificity of insulin action and resistance. Arch. Physiol. Biochem. 117, 96–104 (2011).

3. Pyörälä, M., Miettinen, H., Halonen, P., Laakso, M. & Pyörälä, K. Insulin resistance syndrome predicts the risk of coronary heart disease and stroke in healthy middle-aged men: the 22-year follow-up results of the Helsinki Policemen Study. Arterioscler. Thromb. Vasc. Biol. 20, 538–44 (2000).

4. Brüning, J. C. et al. A muscle-specific insulin receptor knockout exhibits features of the metabolic syndrome of NIDDM without altering glucose tolerance. Mol. Cell 2, 559–69 (1998).

5. Biddinger, S. B. et al. Hepatic insulin resistance is sufficient to produce dyslipidemia and susceptibility to atherosclerosis. Cell Metab. 7, 125–34 (2008).

6. Blüher, M. et al. Adipose tissue selective insulin receptor knockout protects against obesity and obesity-related glucose intolerance. Dev. Cell 3, 25–38 (2002).

7. Gage, M. C. et al. Endothelium-specific insulin resistance leads to accelerated atherosclerosis in areas with disturbed flow patterns: a role for reactive oxygen species. Atherosclerosis 230, 131–9 (2013).

8. Dandona, P. et al. Insulin inhibits intranuclear nuclear factor kappaB and stimulates IkappaB in mononuclear cells in obese subjects: evidence for an anti-inflammatory effect? J. Clin. Endocrinol. Metab. 86, 3257–65 (2001).

9. Dror, E. et al. Postprandial macrophage-derived IL-1β stimulates insulin, and both synergistically promote glucose disposal and inflammation. Nat. Immunol. (2017). doi:10.1038/ni.3659

10. Ding, L. et al. Akt3 deficiency in macrophages promotes foam cell formation and atherosclerosis in mice. Cell Metab. 15, 861–72 (2012).

11. Rotllan, N. et al. Hematopoietic Akt2 deficiency attenuates the progression of atherosclerosis. FASEB J. 29, 597–610 (2015).

12. Babaev, V. R. et al. Macrophage deficiency of Akt2 reduces atherosclerosis in Ldlr null mice. J. Lipid Res. 55, 2296–308 (2014).

13. Iida, K. T. et al. Insulin up-regulates tumor necrosis factor-alpha production in macrophages through an extracellular-regulated kinase-dependent pathway. J. Biol. Chem. 276, 32531–7 (2001).

14. Park, Y. M., R Kashyap, S., A Major, J. & Silverstein, R. L. Insulin promotes macrophage foam cell formation: potential implications in diabetes-related atherosclerosis. Lab. Invest. 92, 1171–80 (2012).

15. Iida, K. T. Insulin Inhibits Apoptosis of Macrophage Cell Line, THP-1 Cells, via Phosphatidylinositol-3-Kinase-Dependent Pathway. Arterioscler. Thromb. Vasc. Biol. 22, 380–386 (2002).

16. Su, D. et al. FoxO1 links insulin resistance to proinflammatory cytokine IL-1beta production in macrophages. Diabetes 58, 2624–33 (2009).

17. Yan, H. et al. Insulin inhibits inflammation and promotes atherosclerotic plaque stability via PI3K-Akt pathway activation. Immunol. Lett. 170, 7–14 (2015).

18. Mauer, J. et al. Myeloid cell-restricted insulin receptor deficiency protects against obesity-induced inflammation and systemic insulin resistance. PLoS Genet. 6, e1000938 (2010).

19. Han, S. et al. Macrophage insulin receptor deficiency increases ER stress-induced apoptosis and necrotic core formation in advanced atherosclerotic lesions. Cell Metab. 3, 257–66 (2006).

20. Baumgartl, J. et al. Myeloid lineage cell-restricted insulin resistance protects apolipoproteinE-deficient mice against atherosclerosis. Cell Metab. 3, 247–56 (2006).

21. Lackey, D. E. & Olefsky, J. M. Regulation of metabolism by the innate immune system. Nat. Rev. Endocrinol. 12, 15–28 (2015).

22. Gage, M. C. et al. Disrupting LXRα phosphorylation promotes FoxM1 expression and modulates atherosclerosis by inducing macrophage proliferation. Proc. Natl. Acad. Sci. 201721245 (2018). doi:10.1073/pnas.1721245115

23. McNelis, J. C. & Olefsky, J. M. Macrophages, Immunity, and Metabolic Disease. Immunity 41, 36–48 (2014).

24. York, A. G. et al. Limiting Cholesterol Biosynthetic Flux Spontaneously Engages Type I IFN Signaling. Cell 163, 1716–1729 (2015).

25. Dang, E. V., McDonald, J. G., Russell, D. W. & Cyster, J. G. Oxysterol Restraint of Cholesterol Synthesis Prevents AIM2 Inflammasome Activation. Cell 171, 1057–1071.e11 (2017).

26. Reboldi, A. et al. 25-Hydroxycholesterol suppresses interleukin-1-driven inflammation downstream of type I interferon. Science (80-.). 345, 679–684 (2014).

27. Gold, E. S. et al. ATF3 protects against atherosclerosis by suppressing 25-hydroxycholesterol-induced lipid body formation. J. Exp. Med. 209, 807–17 (2012).

28. Ley, K., Miller, Y. I. & Hedrick, C. C. Monocyte and macrophage dynamics during atherogenesis. Arterioscler. Thromb. Vasc. Biol. 31, 1506–16 (2011).

29. (CDC)., C. for D. C. Centers for Disease Control & Prevention (CDC) Kaposi’s Sarcoma and Pneumocystis Pneumonia Among Homosexual Men — New York City and Linked references are available on JSTOR for this article : All use subject to JSTOR Terms and Conditions. J. Clin. Invest. 30, 305–308 (1981).

30. Suzuki, R. et al. Diabetes and insulin in regulation of brain cholesterol metabolism. Cell Metab. 12, 567–579 (2010).

31. Clément, S. et al. The lipid phosphatase SHIP2 controls insulin sensitivity. Nature 409, 92–7 (2001).

32. Dubois, E. et al. Developmental defects and rescue from glucose intolerance of a catalytically-inactive novel Ship2 mutant mouse. Cell. Signal. 24, 1971–80 (2012).

33. Wu, X. & Williams, K. J. NOX4 pathway as a source of selective insulin resistance and responsiveness. Arterioscler. Thromb. Vasc. Biol. 32, 1236–45 (2012).

34. Wu, X., Chen, K. & Williams, K. J. The role of pathway-selective insulin resistance and responsiveness in diabetic dyslipoproteinemia. Curr. Opin. Lipidol. 23, 334–44 (2012).

35. Marion, E. et al. The Gene INPPL1, Encoding the Lipid Phosphatase. (2002).

36. Kaisaki, P. J. et al. Polymorphisms in type II SH2 domain-containing inositol 5-phosphatase (INPPL1, SHIP2) are associated with physiological abnormalities of the metabolic syndrome. Diabetes 53, 1900–4 (2004).

37. Marçano, A. C. B. et al. Genetic association analysis of inositol polyphosphate phosphatase-like 1 (INPPL1, SHIP2) variants with essential hypertension. J. Med. Genet. 44, 603–5 (2007).

38. Kagawa, S. et al. Impact of SRC homology 2-containing inositol 5’-phosphatase 2 gene polymorphisms detected in a Japanese population on insulin signaling. J. Clin. Endocrinol. Metab. 90, 2911–9 (2005).

39. Ishida, S. et al. Association of SH-2 containing inositol 5’-phosphatase 2 gene polymorphisms and hyperglycemia. Pancreas 33, 63–7 (2006).

40. Pineda-Torra, I., Gage, M., de Juan, A. & Pello, O. M. Isolation, Culture, and Polarization of Murine Bone Marrow-Derived and Peritoneal Macrophages. Methods Mol. Biol. 1339, 101–9 (2015).

41. Pourcet, B. et al. LXRα regulates macrophage arginase 1 through PU.1 and interferon regulatory factor 8. Circ. Res. 109, 492–501 (2011).

42. Watt, N. T. et al. Endothelial SHIP2 Suppresses Nox2 NADPH Oxidase-Dependent Vascular Oxidative Stress, Endothelial Dysfunction and Systemic Insulin Resistance. Diabetes (2017). at <http://diabetes.diabetesjournals.org/content/early/2017/08/21/db17-0062.long>

43. Strowig, T., Henao-Mejia, J., Elinav, E. & Flavell, R. Inflammasomes in health and disease. Nature 481, 278–86 (2012).

44. Tokarz, V. L., MacDonald, P. E. & Klip, A. The cell biology of systemic insulin function. J. Cell Biol. 217, 2273–2289 (2018).

45. Rask-Madsen, C. & Kahn, C. R. Tissue-specific insulin signaling, metabolic syndrome, and cardiovascular disease. Arterioscler. Thromb. Vasc. Biol. 32, 2052–9 (2012).

46. Horton, J. D., Goldstein, J. L. & Brown, M. S. SREBPs: activators of the complete program of cholesterol and fatty acid synthesis in the liver. J. Clin. Invest. 109, 1125–1131 (2002).

47. Leavens, K., Easton, R., Shulman, G. & Previs, S. Akt2 is required for hepatic lipid accumulation in models of insulin resistance. Cell Metab. 10, 405–418 (2009).

48. Feingold, K. R. et al. Mechanisms of triglyceride accumulation in activated macrophages. J. Leukoc. Biol. 92, 829–39 (2012).

49. Park, K. & Scott, A. L. Cholesterol 25-hydroxylase production by dendritic cells and macrophages is regulated by type I interferons. J. Leukoc. Biol. 88, 1081–1087 (2010).

50. Spann, N. J. et al. Regulated Accumulation of Desmosterol Integrates Macrophage Lipid Metabolism and Inflammatory Responses. Cell 151, 138–152 (2012).

51. Jackson, S. J. et al. Does age matter? The impact of rodent age on study outcomes. Lab. Anim. 51, 160–169 (2017).

52. Senokuchi, T. et al. Forkhead transcription factors (FoxOs) promote apoptosis of insulin-resistant macrophages during cholesterol-induced endoplasmic reticulum stress. Diabetes 57, 2967–76 (2008).

53. Kim, K. et al. Transcriptome Analysis Reveals Nonfoamy Rather Than Foamy Plaque Macrophages Are Proinflammatory in Atherosclerotic Murine Models. Circ. Res. 123, 1127–1142 (2018).

